# Genome-Wide Association for Itraconazole Sensitivity in Non-Resistant Clinical Isolates of *Aspergillus fumigatus*

**DOI:** 10.1101/2020.08.31.275297

**Authors:** Shu Zhao, Wenbo Ge, Akira Watanabe, Jarrod R. Fortwendel, John G. Gibbons

**Author notes:** Corresponding Authors: John G. Gibbons.

## Abstract

*Aspergillus fumigatus* is a potentially lethal opportunistic pathogen that infects over ∼200,000 people and causes ∼100,000 deaths per year globally. Treating *A. fumigatus* infections is particularly challenging because of the recent emergence of azole-resistance. The majority of studies focusing on the molecular mechanisms underlying azole resistance have examined azole-resistant isolates. However, isolates that are susceptible to azoles also display variation in their sensitivity, presenting a unique opportunity to identify genes contributing to azole sensitivity. Here, we used genome-wide association (GWA) analysis to identify loci involved in azole sensitivity by analyzing the association between 68,853 SNPs and itraconazole (ITCZ) minimum inhibitory concentration (MIC) in 76 clinical isolates of *A. fumigatus* from Japan. Population structure analysis suggests the presence of four distinct populations, with ITCZ MICs distributed relatively evenly across populations. We independently conducted GWA when treating ITCZ MIC as a quantitative trait and a binary trait and identified two SNPs with strong associations that were identified in both analyses. These SNPs fell within the coding regions of *Afu2g02220* and *Afu2g02140*. We functionally validated *Afu2g02220* by knocking it out using a CRISPR/Cas-9 approach, because orthologs of this gene are involved in sterol modification and ITCZ targets the ergosterol pathway. Knockout strains displayed no difference in growth compared to the parent strain in minimal media, yet a minor but consistent inhibition of growth in the presence of 0.15 ug/ml ITCZ. Our results suggest that GWA paired with efficient gene deletion is a powerful and unbiased strategy for identifying the genetic basis of complex traits in *A. fumigatus*.

**Importance:** *Aspergillus fumigatus* is a pathogenic mold that can infect and kill individuals with compromised immune systems. The azole class of drugs provide antifungal activity against *A. fumigatus* infections and have become an essential treatment strategy. Unfortunately, *A. fumigatus* azole resistance has recently emerged and rapidly risen in frequency making treatment more challenging. Our understanding of the molecular basis of azole sensitivity has been shaped mainly through candidate gene studies. Unbiased approaches are necessary to understand the full repertoire of genes and genetic variants underlying azole resistance and sensitivity. Here, we provide the first application of genome-wide association analysis in *A. fumigatus* in the identification of a gene (*Afu2g02220*) that contributes to itraconazole susceptibility. Our approach, which combines association mapping and CRISPR/Cas-9 for functional validation of candidate genes, has broad application for investigating the genetic basis of complex traits in fungal systems.

## INTRODUCTION

Fungal infections result in more global deaths per year than deaths from tuberculosis or malaria (1). *Aspergillus fumigatus* is one of the most deadly fungal pathogens and results in more than one hundred thousand deaths per year (1). Invasive aspergillosis (IA) is the most severe infection caused by *A. fumigatus* and occurs when fungal growth, most commonly originating in the lung, disseminates to other parts of the body via the bloodstream (2). *A. fumigatus* is an opportunistic pathogen primarily affecting immunocompromised individuals, and unfortunately, infections have become more common due to the increased usage of immunosuppressive drugs to treat autoimmune disorders and to increase the success of organ transplantation surgery (3-5). Even when aggressively treated with first and second-line antifungal medication, mortality rates can exceed 50% in IA patients (2, 6). The relatively rapid emergence of *A. fumigatus* antifungal resistance has made treatment of infections particularly challenging.

The fungal cell wall is distinct from the mammalian cell wall in several ways and is thus the target of the three most common antifungal drug classes. For example, the echinocandins target β 1,3 glucan, the most abundant polysaccharide in the fungal cell wall, while amphotericin B (a polyene class of antifungal drug) and triazoles (the azole class of antifungal drugs) target ergosterol (7). Ergosterol plays an essential functional role in regulating permeability and fluidity. Triazoles, such as itraconazole (ITCZ) and voriconazole, target the lanosterol demethylase enzymes (Cyp51A and Cyp51B in *A. fumigatus*) which are directly involved in the biosynthesis of ergosterol (8, 9). Blocking Cyp51A and Cyp51B results in the accumulation of a toxic sterol intermediate that causes severe membrane stress, impairment of growth, and cell death(8). Triazoles are the most common first-line treatment for *A. fumigatus* infections.

Strains of *A. fumigatus* have gained resistance to triazoles through mutations in both the coding and regulatory regions of *cyp51A*, and through *cyp51A* independent mechanisms (10). The three most commonly described nonsynonymous mutations in *cyp51A* in azole resistance strains are G54, M220, and G448 (10). Protein structure modeling suggests that these mutations disrupt the binding efficiency of azoles to Cyp51A (11, 12). Increased expression of *cyp51A* through a combination of a promoter region repeat and the L98H point mutation can also confer azole resistance (13). Additionally, several transcription factors (e.g. *srbA*(14), *hapE* (15), *atrR* (16), transporters (e.g. *cdr1B* (17), *atrF* (18), various ABC transporters (19) *etc*.), and other functional groups of genes (e.g. genes involved in calcium signaling, iron balance, signaling pathways, and the Hsp90-calcineurin pathway) have been implicated in azole resistance or susceptibility (20).

The numerous genes identified in azole resistance other than *cyp51A* (10) suggests that additional genes with additive minor effects likely play a role in fine-scale differences in azole sensitivity and resistance. Historically, most genes involved in azole resistance in *A. fumigatus* were discovered through a candidate gene approach (10), or through gene expression differences during exposure to azoles (21, 22). However, candidate gene approaches are biased toward genes and pathways of biological interest. Alternatively, genome-wide association (GWA) studies offer a powerful and versatile approach to identify genetic variants that contribute to complex traits, such as *A. fumigatus* ITCZ sensitivity. In GWA, thousands to millions of high-density genetic variants are tested for a statistical association between each variant and a phenotype of interest (23). Microbial GWAS methods have recently been developed (24-26), but have not yet been applied to study the genetic underpinnings of *A. fumigatus* traits (24-26). However, GWA has been used in other fungal species to identify genes and variants associated with virulence in *Heterobasidion annosum* (27), *Saccharomyces cerevisiae* (28), and *Parastagonospora nodorum* (29), fungal communication in *Neurospora crassa* (30) and aggressiveness in *Fusarium graminearum* (31) and *Zymoseptoria tritici* (32). Here, we hypothesized that GWA could be applied in *A. fumigatus* to identify genes with minor effects on ITCZ sensitivity. We performed GWA in 76 non-resistant clinical isolates of *A. fumigatus* and identified a gene that contributes to fine-scale ITCZ sensitivity. More broadly, we demonstrate that GWA in combination with gene disruption is a useful tool for investigating pathogenicity-associated traits in *A. fumigatus*.

## RESULTS

### Population Structure of Clinical *A. fumigatus* Isolates from Japan

We conducted whole genome sequencing (WGS) for 65 isolates of *A. fumigatus* from Japan and analyzed them in combination with an additional 11 previously sequenced isolates (33). Deduplicated, quality trimmed, and adapter trimmed WGS data of the 76 isolates were used for joint SNP calling with GATK (34) and yielded 206,055 SNPs. To reduce the linkage between adjacent SNPs for population structure analysis, we subsampled SNPs so that they were separated by at least 3.5 kb, which yielded 6,324 SNPs. This subsampled dataset was used for population structure and phylogenetic analysis.

Population structure is a main confounding factor in GWA studies that can lead to false positive associations (35). Therefore, we investigated the population structure of the 76 *A. fumigatus* isolates using the model-based approach implemented in ADMIXTURE (36), as well as a non-model approach where population structure is inferred using discriminant analysis of principal components (DAPC) (37). In ADMIXTURE, cross-validation (CV) error was estimated for each *K* from *K*=1-10. The CV error is calculated by systematically withholding data points, and the lowest value represents the best estimate of the number of ancestral populations (38). Using this approach *K*=4 was the most likely population number **(Figure 1A)**. DAPC uses the Bayesian Information Criterion (BIC) to evaluate the optimal number of clusters (*K*). *K*=4 was also the most likely scenario as evaluated by BIC in DAPC **(Figure 1B)**. Population assignment was highly consistent when the entire SNP set was used, or when subsampled datasets consisting of 6,324 or 756 markers were used to limit linkage between markers **(Figure S1)**. At *K*=4, DAPC assigned the 76 isolates into 4 distinct populations with no admixture, while ADMIXTURE assigned 30 of 76 individuals to more than one population. For population assignment, we placed isolates into their respective population based on their largest membership coefficient. Using this approach, only two isolates, IFM51978 and IFM61610 (**Figure 1C**, indicated by black arrows), were assigned into different populations between the two methods. Phylogenetic network analysis further supports the presence of four main populations and individual population assignment into these populations **(Figure S2)**.

**Figure 1.**
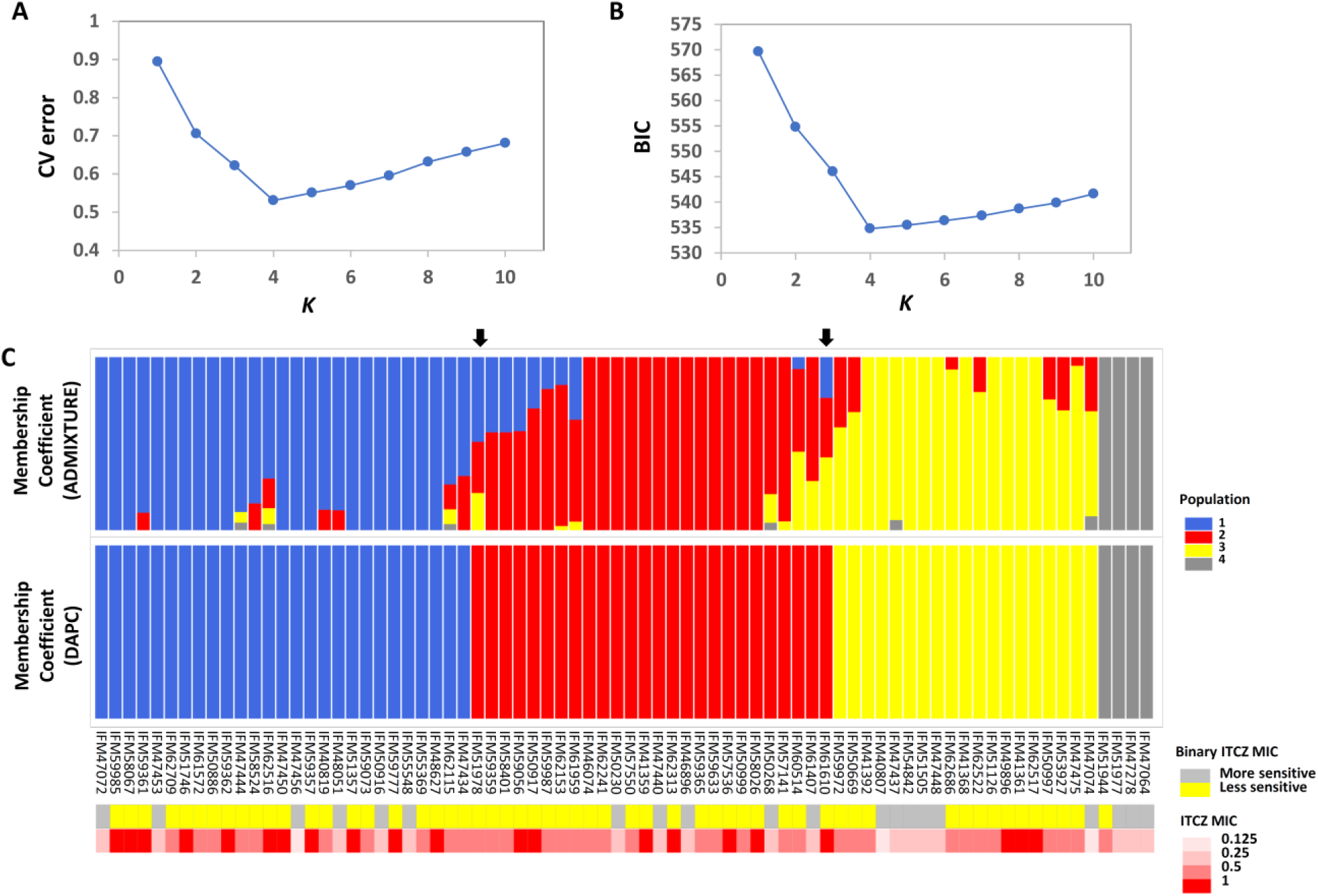
Population structure and ITCZ sensitivity of the 76 Japanese clinical *Aspergillus fumigatus* Isolates. (A) The optimal number of genetic clusters (*K*, X-axis) inferred by ADMIXTURE using cross validation procedure (CV error, Y-axis). (B) The optimal number of genetic clusters (*K*, X-axis) inferred by DAPC using Bayesian Information Criterion (BIC, Y-axis). (C) Membership coefficients (Y-axis) for each of the 76 isolates (X-axis) for ADMIXTURE and DAPC for *K*=4. The two black arrows indicate the two isolates that are assigned into different clusters by ADMIXTURE and DAPC. Population 1, 2, 3 and 4 are colored as blue, red, yellow, and gray, respectively. Binary ITCZ MIC assignment and quantitative ITCZ MIC values are depicted in the upper and lower panels below the membership coefficient plots, respectively. For binary ITCZ MIC, individuals are coded as either more-sensitive (MIC < 0.5, gray) or less-sensitive (MIC ≥ 0.5, yellow).

### Itraconazole Minimum Inhibitory Concentration

The ITCZ MIC of all isolates ranged from 0.125 to 1 ug/ml (MIC_0.125_ = 3, MIC_0.25_ = 17, MIC_0.50_ = 35, and MIC_1_ = 21). For reference, ITCZ resistance is typically defined by MIC ≥ 4 (39). GWA was independently conducted when MIC data was treated as a quantitative trait, and when MIC was treated as a binary trait (“more-sensitive” = MIC < 0.5 or “less-sensitive” = MIC ≥ 0.5). Populations 1, 2, 3, and 4 had 1, 0, 2, and 0 individuals with MIC=0.125, 5, 5, 4, and 3 individuals with MIC=0.25, 10, 14, 10, and 1 individuals with MIC=0.5, and 11, 7, 3, and 0 individuals with MIC=1, respectively **(Figure 1C)**.

### Genome-Wide Association of Itraconazole Sensitivity in *A. fumigatus*

We hypothesized that GWA would allow us to identify genes and/or genetic variants with minor contributions to ITCZ sensitivity. To test this hypothesis, we performed GWA with a set of 68,853 SNPs that have a minor allele frequency >5% and <10% missing data, and the matched ITCZ MICs. Because these isolates have clear population structure **(Figure 1)** we used a mixed effect model GWA, which can reduce the inflated false-positive effect stemming from population structure (25, 40, 41) and has previously been applied in microbial GWA (42, 43). We performed this mixed-model GWA with a covariance matrix as population correction for ITCZ MIC when treated as a quantitative trait **(Figure 2A)** and as a binary trait **(Figure 2B)** using Tassel 5 (44) and RoadTrips (45), respectively. We generated quantile-quantile (Q-Q) plots of expected vs. observed *p-values* to inspect *p-value* inflation, which could be the product of inadequate population structure correction. The Q-Q plots indicate that the distribution of *p-values* for both analyses are not inflated **(Figure S3)**.

**Figure 2.**
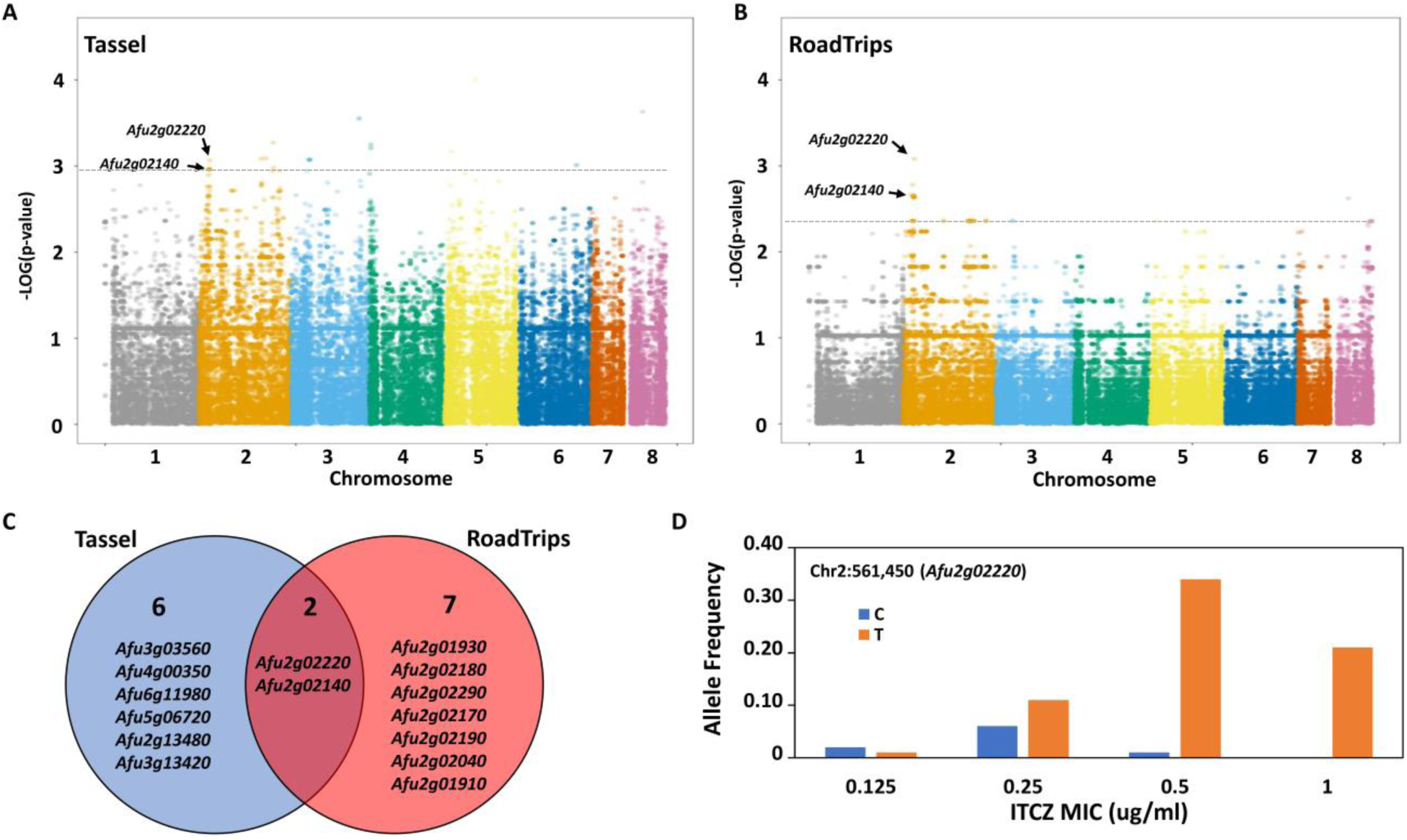
Genome-Wide Association (GWA) for itraconazole (ITCZ) sensitivity. GWA for ITCZ sensitivity when MIC data is treated as a quantitative trait (A) or as a binary trait (B). For binary characterization of ITCZ sensitivity, MIC < 0.5 = more-sensitive and MIC ≥ 0.5 = less-sensitive. The genomic location is of the 68,853 SNPs used for GWA is depicted on the X-axis, while the -log(*P-values*) are depicted on Y-axis. The dotted gray horizontal line represents the cutoff line at the 20^th^ lowest *p-value. Afu2g02140* and *Afu2g02220* were within the 20 SNPs with the strongest associations in both analyses and are labeled on each plot. (C) Venn diagram of the 20 SNPs most strongly associated with ITCZ MIC that overlap genes when data is treated as a quantitative trait (blue circle) and a binary trait (red circle). (D) Allele frequency of the SNP at Chr2:561,450 that falls within *Afu2g02220* (Y-axis) across ITCZ MICs (X-axis).

We considered the 20 SNPs with the lowest *p-values* (lower 0.03 percentile) in each analysis as significant **(Table 1)**. Of the 20 SNPs significantly associated with ITCZ MIC when MIC was treated as a quantitative trait **(Figure 2A)**, 5 SNPs were located in genes (4 in exons and 1 in an intron), 7 SNPs were located in 3’ UTR regions, 2 SNPs were located in 5’ UTR regions, and 6 SNPs were located in intergenic regions **(Table 1)**. Of the four SNPs located in exons, one was synonymous (in *Afu2g02220*) while the remaining three SNPs were non-synonymous (in *Afu2g02140, Afu4g00350*, and *Afu6g11980*) **(Table 1)**. Significant SNPs mapped to chromosomes 2 (N=5), 3 (N=8), 4 (N=2), 5 (N=2), 6 (N=1), and 8 (N=1) **(Figure 2A)**.

**Table 1.**
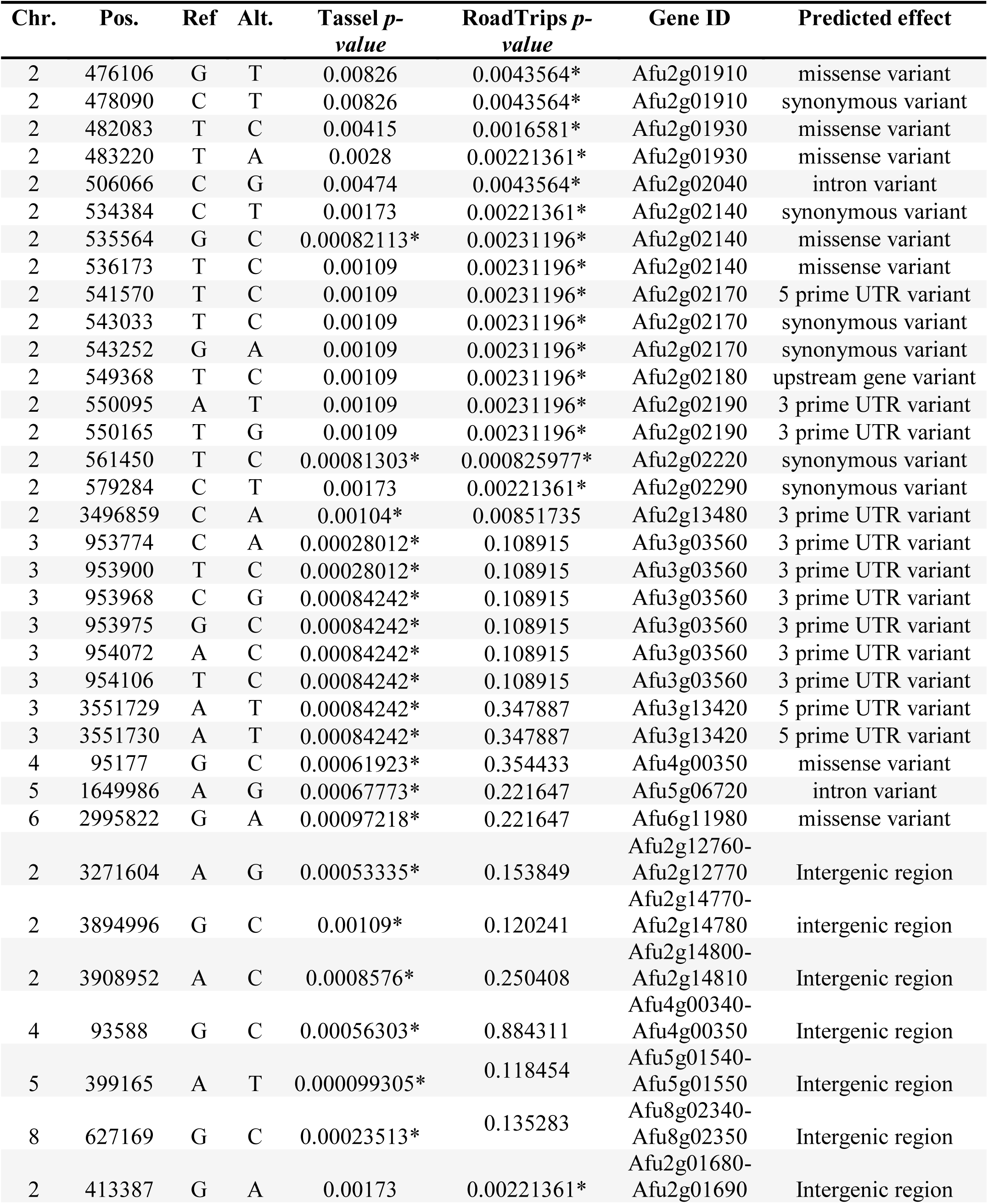
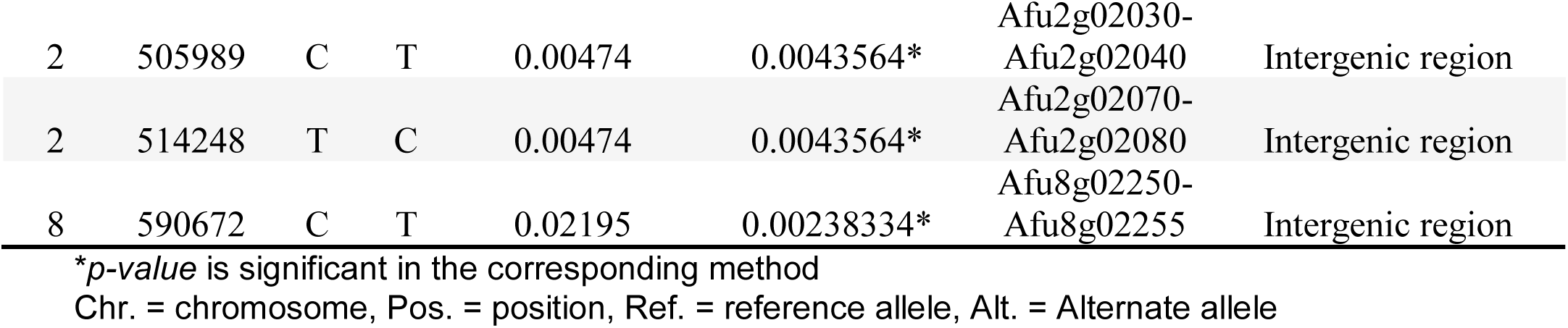
Characterization of SNPs associated with ITCZ sensitivity.

| Chr. | Pos. | Ref | Alt. | Tassel *p*-value | RoadTrips *p*-value | Gene ID | Predicted effect |
| --- | --- | --- | --- | --- | --- | --- | --- |
| 2 | 476106 | G | T | 0.00826 | 0.0043564* | Afu2g01910 | missense variant |
| 2 | 478090 | C | T | 0.00826 | 0.0043564* | Afu2g01910 | synonymous variant |
| 2 | 482083 | T | C | 0.00415 | 0.0016581* | Afu2g01930 | missense variant |
| 2 | 483220 | T | A | 0.0028 | 0.00221361* | Afu2g01930 | missense variant |
| 2 | 506066 | C | G | 0.00474 | 0.0043564* | Afu2g02040 | intron variant |
| 2 | 534384 | C | T | 0.00173 | 0.00221361* | Afu2g02140 | synonymous variant |
| 2 | 535564 | G | C | 0.00082113* | 0.00231196* | Afu2g02140 | missense variant |
| 2 | 536173 | T | C | 0.00109 | 0.00231196* | Afu2g02140 | missense variant |
| 2 | 541570 | T | C | 0.00109 | 0.00231196* | Afu2g02170 | 5 prime UTR variant |
| 2 | 543033 | T | C | 0.00109 | 0.00231196* | Afu2g02170 | synonymous variant |
| 2 | 543252 | G | A | 0.00109 | 0.00231196* | Afu2g02170 | synonymous variant |
| 2 | 549368 | T | C | 0.00109 | 0.00231196* | Afu2g02180 | upstream gene variant |
| 2 | 550095 | A | T | 0.00109 | 0.00231196* | Afu2g02190 | 3 prime UTR variant |
| 2 | 550165 | T | G | 0.00109 | 0.00231196* | Afu2g02190 | 3 prime UTR variant |
| 2 | 561450 | T | C | 0.00081303* | 0.000825977* | Afu2g02220 | synonymous variant |
| 2 | 579284 | C | T | 0.00173 | 0.00221361* | Afu2g02290 | synonymous variant |
| 2 | 3496859 | C | A | 0.00104* | 0.00851735 | Afu2g13480 | 3 prime UTR variant |
| 3 | 953774 | C | A | 0.00028012* | 0.108915 | Afu3g03560 | 3 prime UTR variant |
| 3 | 953900 | T | C | 0.00028012* | 0.108915 | Afu3g03560 | 3 prime UTR variant |
| 3 | 953968 | C | G | 0.00084242* | 0.108915 | Afu3g03560 | 3 prime UTR variant |
| 3 | 953975 | G | C | 0.00084242* | 0.108915 | Afu3g03560 | 3 prime UTR variant |
| 3 | 954072 | A | C | 0.00084242* | 0.108915 | Afu3g03560 | 3 prime UTR variant |
| 3 | 954106 | T | C | 0.00084242* | 0.108915 | Afu3g03560 | 3 prime UTR variant |
| 3 | 3551729 | A | T | 0.00084242* | 0.347887 | Afu3g13420 | 5 prime UTR variant |
| 3 | 3551730 | A | T | 0.00084242* | 0.347887 | Afu3g13420 | 5 prime UTR variant |
| 4 | 95177 | G | C | 0.00061923* | 0.354433 | Afu4g00350 | missense variant |
| 5 | 1649986 | A | G | 0.00067773* | 0.221647 | Afu5g06720 | intron variant |
| 6 | 2995822 | G | A | 0.00097218* | 0.221647 | Afu6g11980 | missense variant |
| 2 | 3271604 | A | G | 0.00053335* | 0.153849 | Afu2g12760-Afu2g12770 | Intergenic region |
| 2 | 3894996 | G | C | 0.00109* | 0.120241 | Afu2g14770-Afu2g14780 | intergenic region |
| 2 | 3908952 | A | C | 0.0008576* | 0.250408 | Afu2g14800-Afu2g14810 | Intergenic region |
| 4 | 93588 | G | C | 0.00056303* | 0.884311 | Afu4g00340-Afu4g00350 | Intergenic region |
| 5 | 399165 | A | T | 0.000099305* | 0.118454 | Afu5g01540-Afu5g01550 | Intergenic region |
| 8 | 627169 | G | C | 0.00023513* | 0.135283 | Afu8g02340-Afu8g02350 | Intergenic region |
| 2 | 413387 | G | A | 0.00173 | 0.00221361* | Afu2g01680-Afu2g01690 | Intergenic region |
| 2 | 505989 | C | T | 0.00474 | 0.0043564* | Afu2g02030-Afu2g02040 | Intergenic region |
| 2 | 514248 | T | C | 0.00474 | 0.0043564* | Afu2g02070-Afu2g02080 | Intergenic region |
| 8 | 590672 | C | T | 0.02195 | 0.00238334* | Afu8g02250-Afu8g02255 | Intergenic region |

\**p-value* is significant in the corresponding method
Chr. = chromosome, Pos. = position, Ref. = reference allele, Alt. = Alternate allele

Of the 20 SNPs significantly associated with ITCZ MIC when MIC was treated as a binary trait **(Figure 2B)**, 12 SNPs were located in genes (11 in exons and 1 in an intron), 2 SNPs were located in 3’ UTR regions, 1 SNP was located in a 5’ UTR regions, and 4 SNPs were located in intergenic regions **(Table 1)**. Of the 11 SNPs located in exons, six were synonymous (in *Afu2g02220, Afu2g02140, Afu2g02290, Afu2g02170* and *Afu2g01910*) while the remaining five were non-synonymous (in *Afu2g01930, Afu2g02140 and Afu2g01910*) **(Table 1)**. Interestingly, in this analysis, 19 of the 20 SNPs with lowest *p-values* were located to a 165 KB region on chromosome 2 (position 413,387 – 579,284) **(Figure 2B)**.

Two significant SNPs overlapped between the quantitative trait and binary trait GWA analyses **(Figure 2C)**. The SNP located in *Afu2g02220* encodes a synonymous variant and had the ninth lowest and lowest *p-values* in the quantitative trait and binary trait analyses, respectively **(Figure 2A, B)**. *Afu2g02220* is annotated as a sterol 3-β-glucosyltransferase **(Table 1)**. The SNP located in *Afu2g02140* encodes a nonsynonymous variant (Ala233Gly) and had the tenth lowest and seventh lowest *p-values* in the quantitative trait and binary trait analyses, respectively **(Figure 2A, B)**. *Afu2g02140* contains a CUE domain (as predicted by PFAM), which has been shown to bind to ubiquitin (46, 47). For both *Afu2g02220* and *Afu2g02140*, the major allele was associated with higher MIC values and the minor allele was absent in all isolates with ITCZ MIC=1, and nearly absent in isolates with ITCZ MIC=0.5 **(Figure 2D, Figure S4)**.

### Expression of *Afu2g02220* and *Afu2g02140* from Existing RNA-seq Experiments

To investigate whether gene expression of *Afu2g02220* and *Afu2g02140* could be modulated by environmental stress, we analyzed *A. fumigatus* RNA-seq data publicly available on FungiDB (48), during oxidative stress, iron depletion, ITCZ exposure, and growth in blood and minimal media (49, 50). *Afu2g02220* was up-regulated during iron starvation (FPKM_control_ = 20.33, FPKM_FeStarvation_ = 32.70, and *p-value* = 5.7e^-4^), oxidative stress induced by H_2_O_2_ (FPKM_control_ = 20.33, FPKM_H2O2_ = 30.61, and *p-value* = 2.7e^-3^), iron starvation + H_2_O_2_ (FPKM_control_ = 20.33, FPKM_FeStarvation+H2O2_ = 39.93, and *p-value* = 6.7e^-23^), and during exposure to ITCZ in strain A1160 (FPKM_-ITCZ_ = 48.20, FPKM_+ITCZ_ = 67.37, and *p-value* = 1.7e^-4^) **(Figure S5A)**. *Afu2g02140* was not significantly up-regulated during any condition, and expressed at lower levels across all conditions compared to *Afu2g02220* **(Figure S5)**.

### Validation of a GWA Candidate Gene via CRISPR/Cas9 Gene Deletion

We chose to functionally examine the role of *Afu2g02220* because (i) the SNP located in this gene had highly significant *p-values* in both GWA analyses (ii) *Afu2g02220* has a predicted functional role in sterol metabolism, and ITCZ targets the ergosterol pathway and (iii) *Afu2g02220* was up-regulated during ITCZ exposure **(Figure S5A)**. Thus, we used an established CRISPR/Cas-9 method (51) to knockout (KO) *Afu2g02220* by replacing it with the indicator gene hygromycin B phosphotransferase *(hygR)* in the *A. fumigatus* CEA10 genetic background **(Figure 3A)**. We generated two independent KOs of *Afu2g02220* which we validated by via PCR **(Figure 3B)**.

**Figure 3.**
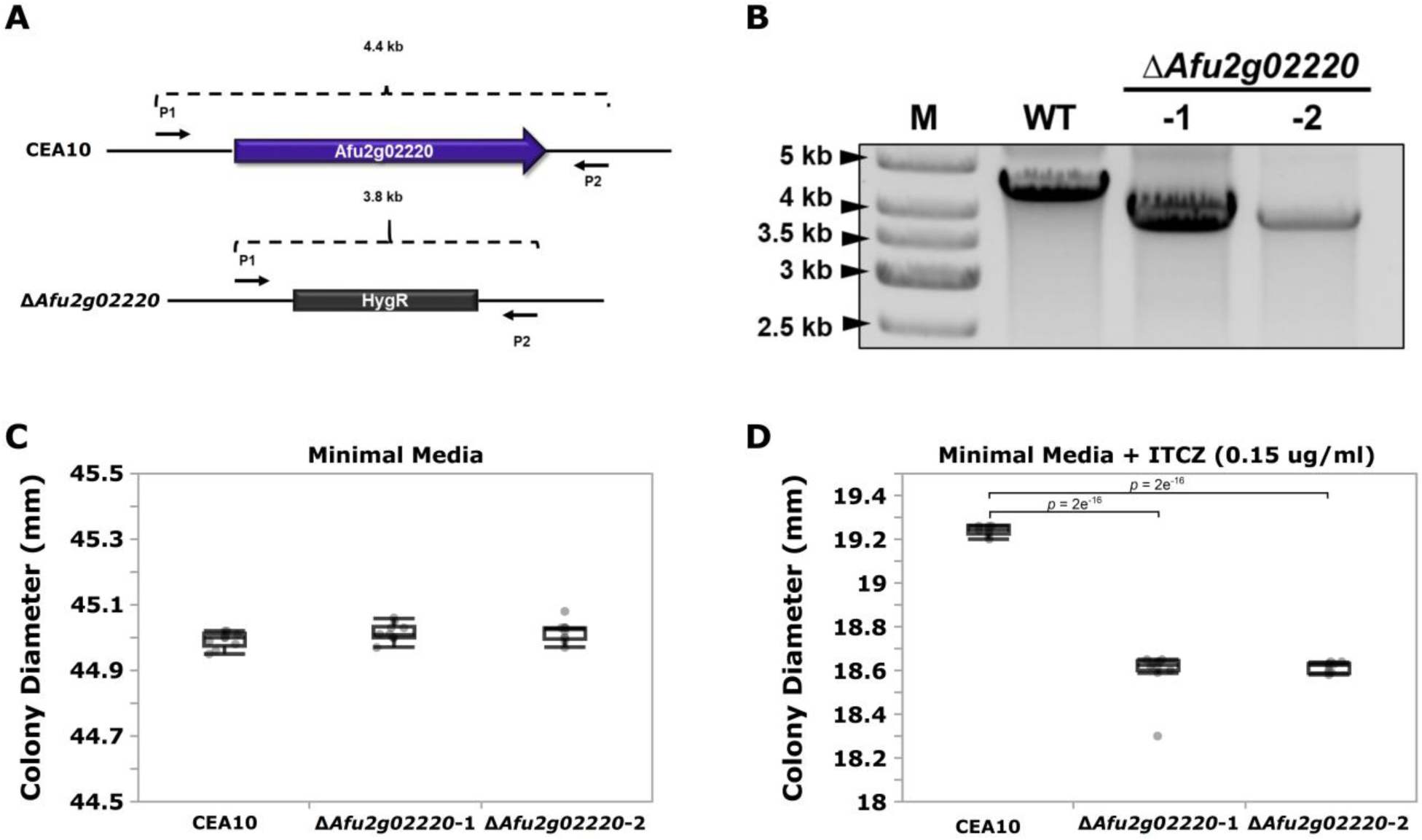
Deletion of *Afu2g02220* impairs growth in the presence of itraconazole (ITCZ). (A) Schematic of *Afu2g02220* gene deletion via CRISPR/Cas-9. The blue box with arrow in the upper panel represents *Afu2g02220* in the parent CEA10 genome (wild type, WT), while the gray box in the lower panel represents the indicator gene *HygR* that replaced *Afu2g02220* in *ΔAfu2g02220* strains. The two black arrows on the flanking region of the locus indicate the forward primer (P1) and reverse primer (P2) used for PCR validation. The WT amplicon size is ∼4.4 Kb, while the *HygR* gene replacement amplicon is ∼3.8 Kb. (B) Validation of *Afu2g02220* gene replacement via PCR. Lanes “M”, “WT”, “-1” and “-2” indicate ladder, PCR product from WT and PCR product from the two independent knockout strains, respectively. Boxplots for colony diameter of WT and *ΔAfu2g02220* strains grown at 37°C for 72 hours on minimal media (C) and minimal media with 0.15 ug/ml ITCZ (D). Measurements were collected for 10 biological replicates for each experiment. Dunnett’s test *p-values* indicate a significant reduction in growth in the KOs compared to the WT.

To test the effect of *Afu2g02220* on ITCZ sensitivity, we grew the wild type (WT) and Δ*Afu2g02220* strains in the presence of 0.15 ug/ml of ITCZ and measured colony diameter after 72 hours of incubation at 37°C. We observed a qualitative reduction in conidia production in KO strains **(Figure S6)**. In minimal media without ITCZ Δ*Afu2g02220-1* and Δ*Afu2g02220-2* growth rates did not significantly differ from the WT (Δ*Afu2g02220-1* = 45.016 mm, Δ*Afu2g02220-2* = 45.018 mm, WT = 44.994 mm) **(Figure 3C)**. This result suggests that the background growth rate of Δ*Afu2g02220* is not impacted by the gene deletion. However, at ITCZ concentrations of 0.15 ug/ml we observed a minor but consistent reduction in growth in KO strains compared to WT (Δ*Afu2g02220-1* = 18.594 mm, Δ*Afu2g02220-1* = 18.615 mm, WT = 19.239 mm) (*p-value* = 2e^-16^ for both KOs) **(Figure 3D)**. These results suggest that *Afu2g02220* plays a minor role in ITCZ sensitivity.

## DISCUSSION

Here, we analyzed the association between SNP allele frequency and ITCZ MIC data from 76 Japanese clinical isolates of *A. fumigatus* to identify loci involved in ITCZ sensitivity. MIC values fell within a relatively tight range of 0.125 to 1 ug/ml (for reference, ITCZ resistant strains are defined by MICs ≥ 4 ug/ml (39)). We reasoned that GWA could be a feasible tool to identify loci that contribute to the small differences in ITCZ MIC we observed across these clinical isolates. We identified several candidate SNPs and loci associated with ITCZ sensitivity, and validated the function of the top candidate by knocking it out using a CRISPR/Cas-9 based approach.

We identified a synonymous variant in *Afu2g02220* that showed highly significant associations with ITCZ sensitivity across GWA analyses with different underlying statistical models **(Figure 2)**. Synonymous mutations can be functional through their (i) effect on cis-regulatory regions (*e*.*g*. splice sites or miRNA and exonic transcription factor binding sites), (ii) alteration of mRNA structure, or (iii) influence on translation speed (*e*.*g*. codon usage) (52). Determining the mechanism by which this variant alters phenotype would require extensive *in silico* and *in vitro* experimentation. *Afu2g02220* encodes a predicted sterol glycosyltransferase. This enzyme biosynthesizes sterol glucosides, which make up the common eukaryotic membrane bound lipids. Orthologs of *Afu2g02220* from the ascomycete yeasts *Saccharomyces cerevisiae (Atg26), Candida albicans, Pichia pastoris*, as well as the amoeba *Dictyostelium discoideum* can use various sterols, including ergosterol, as sugar acceptors (53). In *S. cerevisiae*, Atg26 can directly bind to and glycosylate ergosterol, which yields ergosterol-glucoside (54). In *S. cerevisiae* Δ*Atg26* did not impair growth when cultured in complex or minimal media, low or elevated temperatures, varying osmotic stress conditions, or in the presence of nystatin, an antifungal drug that binds to ergosterol (53). Similarly, we did not observe a difference in growth rate between Δ*Afu2g02220* and the WT when grown in minimal media **(Figure 3C)**.

In addition to its role in sterol modification, *Afu2g02220* may also have additional functions related to autophagy (55). Orthologs of *Afu2g02220* in *Pichia pastoris* (*PpAtg26*) (56), *Colletotrichum orbiculare* (*CoAtg26*) (57) and *Aspergillus oryzae* (*AoAtg26*) (55) are required for autophagy. In *A. oryzae, ΔAoAtg26* shows deficiency in degradation of peroxisomes, mitochondria, and nuclei and localizes to vacuoles (55). *ΔAoAtg26* also shows reductions in conidiation and impairment of aerial hyphae formation (55). Similarly, we observed a reduction in condition in Δ*Afu2g02220* compared to the WT **(Figure S6)**.

The fungal cell wall is rigid but also dynamic in order to respond to environmental stress. Because Afu2g02220 may directly interact with ergosterol, we hypothesized that environmental stress could alter the expression of *Afu2g02220*. We analyzed *A. fumigatus* RNA-seq data during growth under during iron depletion, oxidative stress, ITCZ exposure and growth in blood and minimal media (48). We found that *Afu2g02220* expression was significantly up-regulated during oxidative stress, iron depletion and ITCZ exposure **(Figure S5)**. However, other studies examining gene expression (21, 22) or protein abundance (58) during exposure to ITCZ and voriconazole (22) (another triazole with the same mechanism of action as ITCZ) did not observe differential abundance of the Afu2g02220 transcript or protein. Additional experiments are necessary to determine the precise role of *Afu2g02220* in stress response and ITCZ sensitivity.

Previously, Palma-Guerrero et al. (2016) used a similar approach to identify NCU04379 as a gene that contributes to fungal communication in *N. crassa*. This study used RNA-seq data to identify genetic variants, Fisher’s exact tests to perform GWA in a closely related group of 112 isolates, and existing deletion mutants generated by the *Neurospora* Genome Project (59, 60) to validate the involvement of NCU04379 in cellular communication during germling fusion. A study in *S. cerevisiae* used a mixed linear model to identify correlations between genotype and tolerance to hydrolysate toxins, and used homologous recombination to knockout candidate genes in two independent genetic backgrounds (61). Interestingly, eight of 14 gene knockouts had a significant effect on phenotype in one, but not both genetic backgrounds, suggesting that the network of genes contributing to hydrolysate toxins tolerance likely differs between genetic backgrounds. The results of these studies, and of our own, broadly suggest that GWA in combination with an efficient gene disruption technique is a powerful and unbiased approach for identifying the genetic basis of polygenic phenotypes in fungal systems.

## MATERIALS AND METHODS

### Japanese *Aspergillus fumigatus* Clinical isolates

Sixty-five *A. fumigatus* clinical strains were provided through the National Bio-Resource Project (NBRP), Japan (http://nbrp.jp/) **(Table S1)**. Whole genome paired-end Illumina sequence data for an additional 11 *A. fumigatus* isolates that were previously sequenced and have ITCZ MIC data (33) **(Table S1)** were downloaded from NCBI Sequence Read Archive (SRA) (62) using the SRA toolkit (https://trace.ncbi.nlm.nih.gov/Traces/sra/sra.cgi?cmd=show&f=software&m=software&s=software).

### Minimum Inhibitory Concentration Testing

Minimal inhibitory concentration (MIC) of ITCZ for each isolate was determined following the Clinical and Laboratory Standards Institute (CLSI) M38-A2 (63) broth microdilution method. Briefly, each strain was incubated in standard RPMI 1640 broth (pH=7) (Sigma Aldrich, St. Louis, US-MO) with a range of ITCZ concentrations at 35°C for 48h. MIC values represent the lowest ITCZ concentrations that completely inhibited growth.

### DNA extraction and Illumina Whole-Genome Sequencing

Genomic DNA (gDNA) isolation was performed as previously described(64). gDNA was directly isolated from conidia stocks using the MasterPure(tm) Yeast DNA Purification Kit (Lucigen/Epicentre) following the manufacturer’s instructions, with several minor modifications. Conidia stocks were centrifuged at 14,000 RPM for 5 minutes to obtain a pellet. Next, 300 ml of yeast cell lysis solution was added to the pellet along with 0.4 ml of sterile 1.0 μm diameter silica beads. Lysis was carried out on a Biospec Mini-BeadBeater-8 at medium intensity for 8 minutes. One ul of RNase was added to the cell lysis solution and incubated at 65° C for 30 minutes. DNA isolation and purification were conducted according to the manufacturer’s instructions for the remainder of the protocol. PCR-free 150-bp paired-end libraries were constructed and sequenced by Novogene (https://en.novogene.com/) on an Illumina NovaSeq 6000.

### Data Availability

Raw whole-genome Illumina data for the 65 isolates are available through NCBI BioProject PRJNA638646 and the 11 previously sequenced isolates by Takahashi-Nakaguchi *et al*. (2015) through NCBI BioProject PRJDB1541.

### Quality Control and Sequence Read Mapping

Raw reads were first deduplicated using tally (65) with the parameters “--with-quality” and “--pair-by-offset” to remove potential PCR duplication during library construction. Next, we used trim_galore v0.4.2 (http://www.bioinformatics.babraham.ac.uk/projects/trim_galore/) to trim residual adapter sequences from reads, and trim reads where quality score was below 30, with the parameters “--stringency 5” and “-q 30” respectively. Trimmed reads shorter than 50 bp were then discarded using the option “--length 50”. Next, the deduplicated and trimmed read set was mapped to the *A fumigatus* Af293 reference genome (66) using BWA-MEM v0.7.15 aligner (67). The resulting SAM files were converted into sorted BAM files using the “view” and “sort” functions in samtools 1.4.1(68).

### SNP genotyping

Because *A. fumigatus* is haploid, we followed the best practice pipeline for “Germline short variant discovery” (69) in Genome Analysis ToolKit (GATK) v4.0.6.0 (34). The function “HaplotypeCaller” was used to call short variants (SNPs and INDELs) with the sorted BAM file for each sample. The resulting g.vcf files of all 76 samples were then combined to generate a joint-called variant file using the function “GenotypeGVCFs”. Next only SNPs were extracted from the joint-called variant file using the function “SelectVariants”. To limit false positive variant calling, the function “VariantFiltration” was used to carry out “hard filtering” with the following parameters: “QD < 25.0 || FS > 5.0 || MQ < 55.0 || MQRankSum < -0.5 || ReadPosRankSum < -2.0 || SOR > 2.5”. 206,055 polymorphic loci were predicted after hard filtering.

### Population Structure of *A. fumigatus* isolates

To investigate the population structure of the *A. fumigatus* isolates we used a subset of population genetic informative SNPs. We used VCFtools v0.1.14 (70) (http://vcftools.sourceforge.net/) with options “--maf 0.05 --max-missing 1 --thin 3500”, to filter the full set of SNPs and require a minor allele frequency ≥ 5%, no missing data across all samples, and at least 3.5 Kb distance between SNPs. 6,324 SNPs remained after filtering, and subsequent population structure analysis was conducted with this marker set. In addition, to test the consistency of population assignments with different number of SNPs, population structure analysis was conducted with a dense SNP set where thinning was not applied (59,433 SNP sites) and an additional thinned SNP set where markers were spaced apart by at least 35 Kb (756 SNPs).

To conduct population structure analysis, we first used the model-based program ADMIXTURE v1.3 (36) for *K*=1-10, where *K* indicates the number of populations. The 5-fold cross-validation (CV) procedure was calculated to find the most likely *K* with option “--cv=5”. For each *K* the CV error was calculated and the *K* with lowest CV error indicated the most likely population number. Additionally, we used the non-model based population structure software DAPC (37) in the adegenet” package v2.1.2 (71) in R v3.5.3 (72) to the predict the number and assignment of individuals into populations. DAPC applies a Bayesian clustering method to identify populations without evolutionary models. The most likely number of populations was inferred by calculating the Bayesian Information Criterion (BIC) for each *K*.

Lastly, we also constructed a phylogenetic network with the alignment of 6,324 SNPs. The phylogenetic network was built using SplitsTree v4.14.4(73) with the neighbor joining method and 1,000 replicates for bootstrap analysis.

### Genome Wide Association Analysis for Itraconazole Sensitivity

Genome Wide Association (GWA) analysis was conducted to identify genetic variants that were significantly correlated with ITCZ MIC. For GWA analysis, we filtered our complete set of SNPs with VCFtools to include SNPs with a minor allele frequency ≥5%, SNP sites with ≤10% missing data, and SNPs that were biallelic. This filtering procedure resulted in 68,853 SNPs were used for GWA.

Two models were used to perform GWA between each of the 68,853 SNPs and ITCZ MIC. When ITCZ MIC data was treated as a quantitative trait (S1. table), we used a linear mixed model with a genetic distance matrix for population structure correction in Tassel(44). GWA was also performed when ITCZ MIC was treated as a binary trait (MIC ≤ 0.5 = more susceptible, and MIC > 0.5 = less susceptible). In this GWA analysis, we used a mixed effect logistic model with an empirical covariance matrix as a population structure correction in RoadTrips(45). Quantile– quantile(Q-Q) plots were generated using the R package “qqman” (74) in order to evaluate potential *p-value* inflation. The potential functional effects of candidate SNPs were predicted using SnpEff v4.3t (75) with the *A. fumigatus* Af293 reference genome annotation.

### RNA-seq based Expression data from *Afu2g02220* and *Afu2g02140*

To investigate the expression patterns of our candidate genes *Afu2g02220* and *Afu2g02140*, we obtained FPKM values as well as fold-change and *p-values* for pairwise comparisons from FungiDB (https://fungidb.org/fungidb/) (48) for oxidative stress, iron depletion, growth in blood and minimal media, and ITCZ exposure (49, 50).

### Gene deletion of *Afu2g02220* in *A. fumigatus* CEA10

*A. fumigatus* strain CEA10 was used as the genetic background for the deletion of *Afu2g02220*. The deletion was carried out using a clustered regularly interspaced short palindromic repeats (CRISPR)/Cas9-mediated protocol for gene editing, as previously described (51). Briefly, two Protospacer Adjacent Motif (PAM) sites, at both upstream and downstream of *Afu2g02220*, were selected using the EuPaGDT tool (76) and custom crRNAs were designed using the 20 base pairs of sequence immediately upstream of the PAM site. The crRNA used are as follows: 5’ crRNA of *Afu2g0222*0 = CTGTTATTTTCTTCGGGTCT and 3’ crRNA of *Afu2g02220* = TGGACCAGGAAGAAACTGAG. Both crRNAs were purchased form a commercial vendor. Complete guideRNAs (gRNAs) were then assembled *in vitro* using the custom designed crRNA coupled with a commercially acquired tracrRNA. The assembled gRNAs were then combined with commercially purchased Cas9 to form ribonucleoproteins for transformation, as previously described (51). Repair templates carrying a hygromycin resistance (HygR) cassette were PCR amplified to contain 40-basepair regions of microhomology on either side for homologous integration at the double strand DNA break induced by the Cas9 nuclease. Protoplast-mediated transformations were then carried using the hygromycin repair templates and Cas-ribonucleoproteins for gene targeting. Homologous integrations were confirmed by PCR. The primers used are as follows:

*Afu2g02020* KO Forward Screening Primer (P1): GGATGCGTTGTTCCTGTGCG *Afu2g02220* KO Reverse Screening Primer (P2): AACGAGGGCTGGAGTGCC Common *HygR* Reverse Screening Primer (P3): ACACCCAATACGCCGGCC

### Fungal Growth in Presence of ITCZ

Conidia (10^4^) were inoculated onto GMM agar (77), and GMM agar supplemented with 0.15 ug/ml ITCZ. Plates were incubated for 72 hours at 37°C. As a measure of drug sensitivity, colony diameter was measured with a digital caliper for the ***Δ****Afu2g02220* strains and the CEA10 parent strain. Experiments were performed with ten replicates.

## ACKNOWLEGDMENTS

This research was supported by grant 1R21AI137485-01 from the National Institutes of Health and National Institutes of Allergy and Infectious Diseases to JGG which supports JGG and SZ. JRF and WG are supported by NIAID R01AI143197 to JRF. AW was supported by the Japan Agency for Medical Research and Development (AMED) under Grant Numbers 20jm0110015.

---

**Table S1.** Sample information for the 76 Japanese clinical *A. fumigatus* isolates.

| Sample_ID | ITCZ_MIC<br>(ug/ml) | ITCZ_MIC<br>(binary) | BioProject<br>Accession<br>Number | BioSample<br>Accession<br>Number |
| --- | --- | --- | --- | --- |
| IFM58026 | 1 | 2 | PRJDB1541 | SAMD00013550 |
| IFM58401 | 0.5 | 2 | PRJDB1541 | SAMD00013548 |
| IFM59056 | 1 | 2 | PRJDB1541 | SAMD00013552 |
| IFM59361 | 1 | 2 | PRJDB1541 | SAMD00013551 |
| IFM59365 | 0.5 | 2 | PRJDB1541 | SAMD00013555 |
| IFM59777 | 1 | 2 | PRJDB1541 | SAMD00013547 |
| IFM60514 | 0.5 | 2 | PRJDB1541 | SAMD00013544 |
| IFM61407 | 0.25 | 1 | PRJDB1541 | SAMD00013549 |
| IFM61610 | 1 | 2 | PRJDB1541 | SAMD00013558 |
| IFM62115 | 0.5 | 2 | PRJDB1541 | SAMD00013556 |
| IFM62516 | 1 | 2 | PRJDB1541 | SAMD00013559 |
| IFM40807 | 0.125 | 1 | PRJNA638646 | SAMN15199827 |
| IFM40819 | 0.5 | 2 | PRJNA638646 | SAMN15199828 |
| IFM41359 | 1 | 2 | PRJNA638646 | SAMN15199829 |
| IFM41361 | 1 | 2 | PRJNA638646 | SAMN15199830 |
| IFM41368 | 0.5 | 2 | PRJNA638646 | SAMN15199831 |
| IFM41392 | 0.5 | 2 | PRJNA638646 | SAMN15199832 |
| IFM46074 | 0.5 | 2 | PRJNA638646 | SAMN15199833 |
| IFM46896 | 0.25 | 1 | PRJNA638646 | SAMN15199834 |
| IFM47064 | 0.25 | 1 | PRJNA638646 | SAMN15199835 |
| IFM47072 | 0.25 | 1 | PRJNA638646 | SAMN15199836 |
| IFM47074 | 0.125 | 1 | PRJNA638646 | SAMN15199837 |
| IFM47278 | 0.25 | 1 | PRJNA638646 | SAMN15199838 |
| IFM47434 | 0.5 | 2 | PRJNA638646 | SAMN15486847 |
| IFM47437 | 0.25 | 1 | PRJNA638646 | SAMN15199839 |
| IFM47440 | 0.25 | 1 | PRJNA638646 | SAMN15199840 |
| IFM47444 | 0.5 | 2 | PRJNA638646 | SAMN15199841 |
| IFM47448 | 0.25 | 1 | PRJNA638646 | SAMN15486848 |
| IFM47450 | 1 | 2 | PRJNA638646 | SAMN15199842 |
| IFM47453 | 0.25 | 1 | PRJNA638646 | SAMN15199843 |
| IFM47456 | 0.125 | 1 | PRJNA638646 | SAMN15199844 |
| IFM47475 | 0.5 | 2 | PRJNA638646 | SAMN15199845 |
| IFM48051 | 0.25 | 1 | PRJNA638646 | SAMN15199846 |
| IFM48627 | 1 | 2 | PRJNA638646 | SAMN15486849 |
| IFM49896 | 1 | 2 | PRJNA638646 | SAMN15199847 |
| IFM50230 | 0.25 | 1 | PRJNA638646 | SAMN15199848 |
| IFM50268 | 0.25 | 1 | PRJNA638646 | SAMN15199849 |
| IFM50669 | 0.5 | 2 | PRJNA638646 | SAMN15199850 |
| IFM50886 | 0.5 | 2 | PRJNA638646 | SAMN15486850 |
| IFM50916 | 0.25 | 1 | PRJNA638646 | SAMN15199851 |
| IFM50917 | 1 | 2 | PRJNA638646 | SAMN15199852 |
| IFM50997 | 0.5 | 2 | PRJNA638646 | SAMN15199853 |
| IFM50999 | 0.5 | 2 | PRJNA638646 | SAMN15199854 |
| IFM51126 | 0.5 | 2 | PRJNA638646 | SAMN15199855 |
| IFM51357 | 1 | 2 | PRJNA638646 | SAMN15199856 |
| IFM51505 | 0.25 | 1 | PRJNA638646 | SAMN15199857 |
| IFM51746 | 1 | 2 | PRJNA638646 | SAMN15199858 |
| IFM51944 | 0.5 | 2 | PRJNA638646 | SAMN15199859 |
| IFM51977 | 0.25 | 1 | PRJNA638646 | SAMN15199860 |
| IFM51978 | 0.5 | 2 | PRJNA638646 | SAMN15199861 |
| IFM53927 | 0.5 | 2 | PRJNA638646 | SAMN15199862 |
| IFM54842 | 0.25 | 1 | PRJNA638646 | SAMN15199863 |
| IFM55369 | 0.5 | 2 | PRJNA638646 | SAMN15199864 |
| IFM55548 | 0.25 | 1 | PRJNA638646 | SAMN15486852 |
| IFM57141 | 0.5 | 2 | PRJNA638646 | SAMN15199865 |
| IFM57536 | 1 | 2 | PRJNA638646 | SAMN15199866 |
| IFM57550 | 0.5 | 2 | PRJNA638646 | SAMN15199867 |
| IFM58067 | 1 | 2 | PRJNA638646 | SAMN15199868 |
| IFM58524 | 0.5 | 2 | PRJNA638646 | SAMN15199869 |
| IFM59073 | 0.5 | 2 | PRJNA638646 | SAMN15199870 |
| IFM59357 | 1 | 2 | PRJNA638646 | SAMN15199871 |
| IFM59359 | 0.5 | 2 | PRJNA638646 | SAMN15199872 |
| IFM59362 | 1 | 2 | PRJNA638646 | SAMN15199873 |
| IFM59633 | 0.5 | 2 | PRJNA638646 | SAMN15199874 |
| IFM59972 | 0.5 | 2 | PRJNA638646 | SAMN15199875 |
| IFM59985 | 1 | 2 | PRJNA638646 | SAMN15199876 |
| IFM59987 | 0.5 | 2 | PRJNA638646 | SAMN15199877 |
| IFM61572 | 0.5 | 2 | PRJNA638646 | SAMN15199878 |
| IFM61959 | 0.5 | 2 | PRJNA638646 | SAMN15199879 |
| IFM62153 | 0.5 | 2 | PRJNA638646 | SAMN15199880 |
| IFM62241 | 0.5 | 2 | PRJNA638646 | SAMN15486855 |
| IFM62313 | 1 | 2 | PRJNA638646 | SAMN15199881 |
| IFM62517 | 1 | 2 | PRJNA638646 | SAMN15199882 |
| IFM62522 | 0.5 | 2 | PRJNA638646 | SAMN15486856 |
| IFM62686 | 0.5 | 2 | PRJNA638646 | SAMN15199883 |
| IFM62709 | 0.5 | 2 | PRJNA638646 | SAMN15199884 |

\*Binary MIC 1 = ITCZ MIC < 0.5 ug/ml and 2 = ITCZ MIC ≥ 0.5 ug/ml

**Figure S1.**
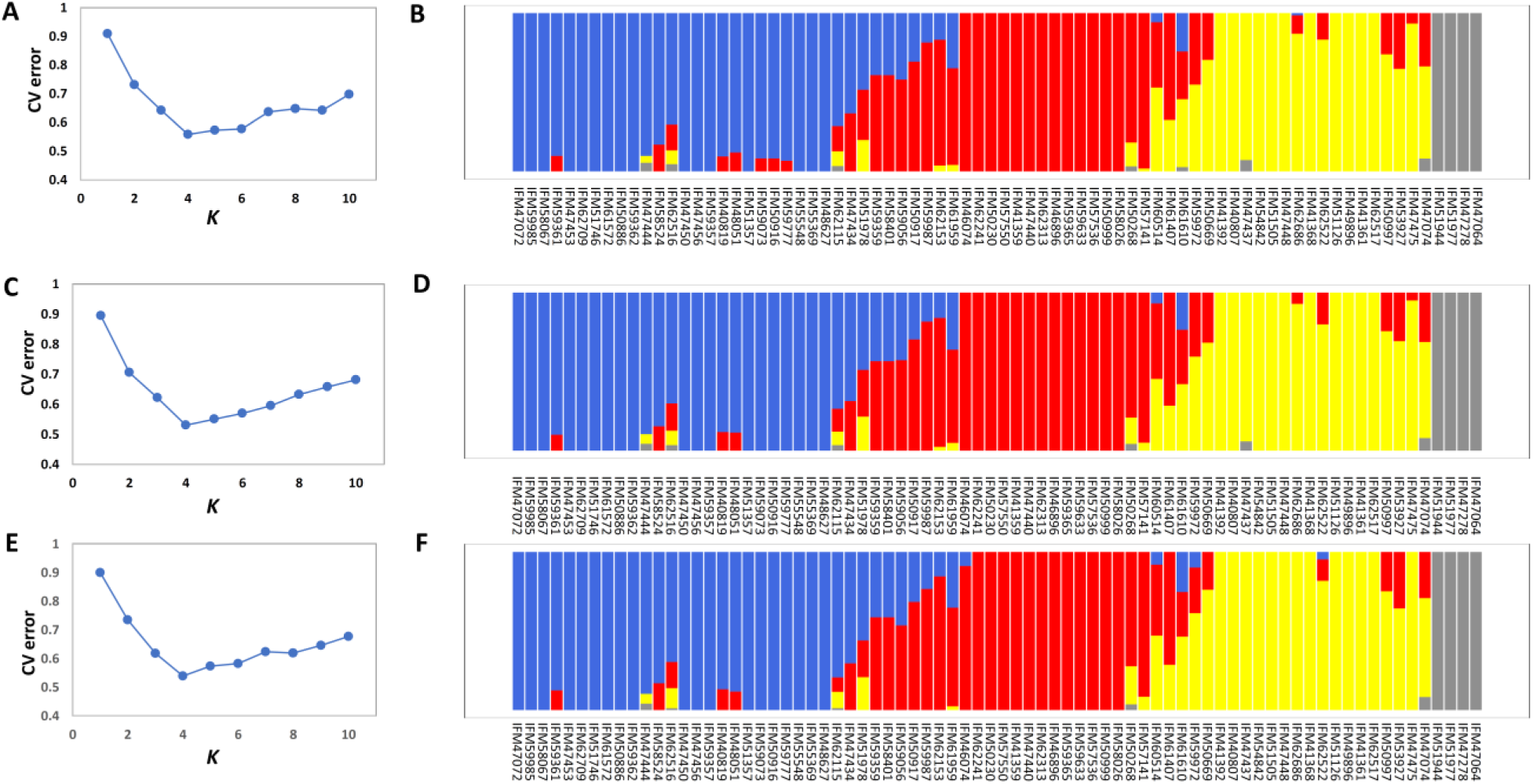
ADMIXTURE based estimate of the optimal predicted population number and population assignment. The optimal predicted population number (*K*) (X-axis) estimated with the CV error (Y-axis) and membership coefficient plots for *K*=4 using 59,433 SNPs (A, B), 6,884 SNPs (with at least 3.5 kb distance between SNPs) (C, D), and 756 SNPs (with at least 35 kb distance between SNPs) (E, F).

**Figure S2.**
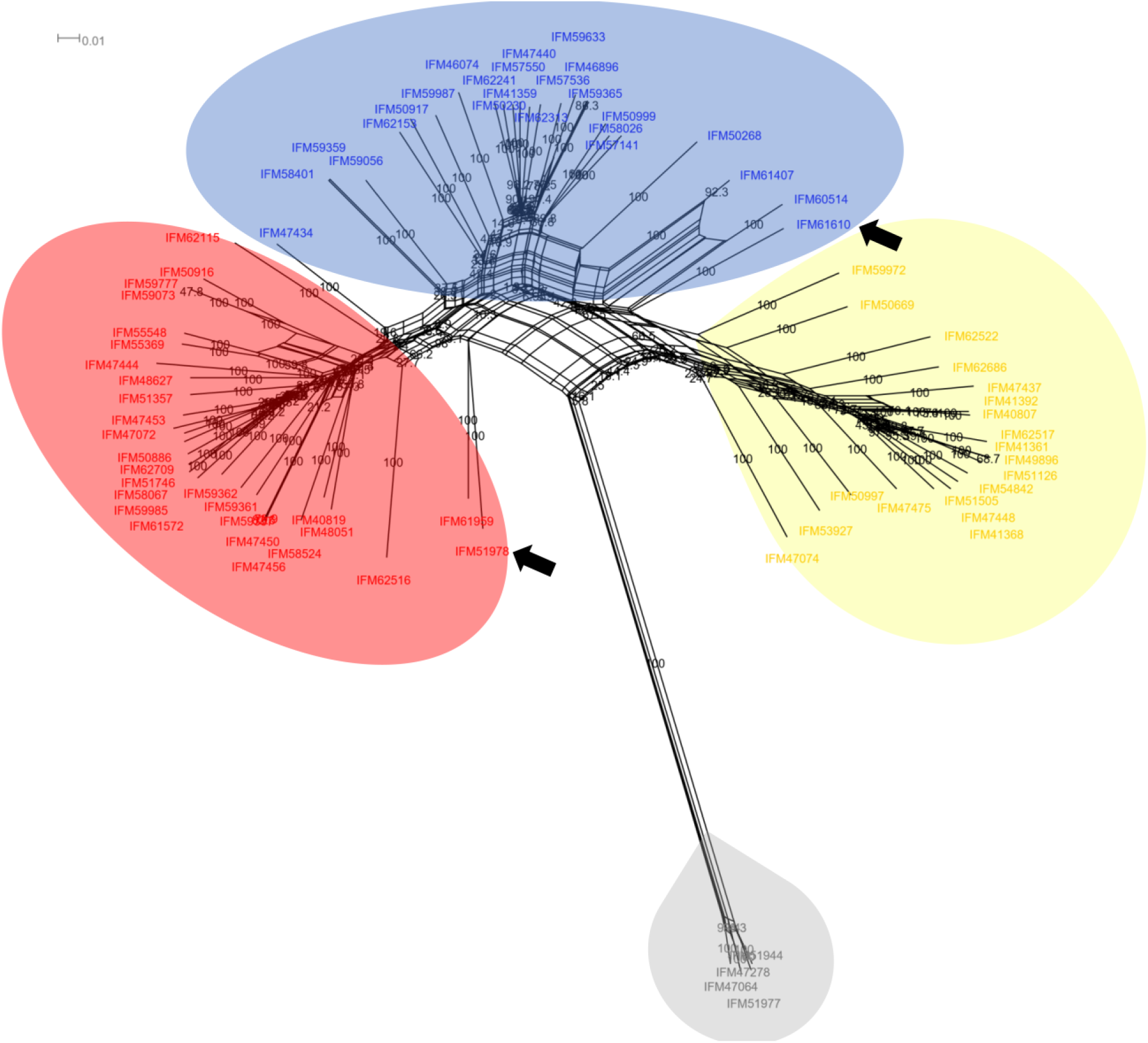
Phylogenetic network of the 76 *A. fumigatus* Japanese clinical isolates. The scale bar represents the proportion of nucleotide differences between two isolates. Isolates that are assigned to the DAPC based populations 1, 2, 3 and 4 are colored as blue, red, yellow, and gray, respectively. The two isolates that are assigned into different clusters by ADMIXTURE and DAPC are indicated with black arrows. Bootstrap values are indicated.

**Figure S3.**
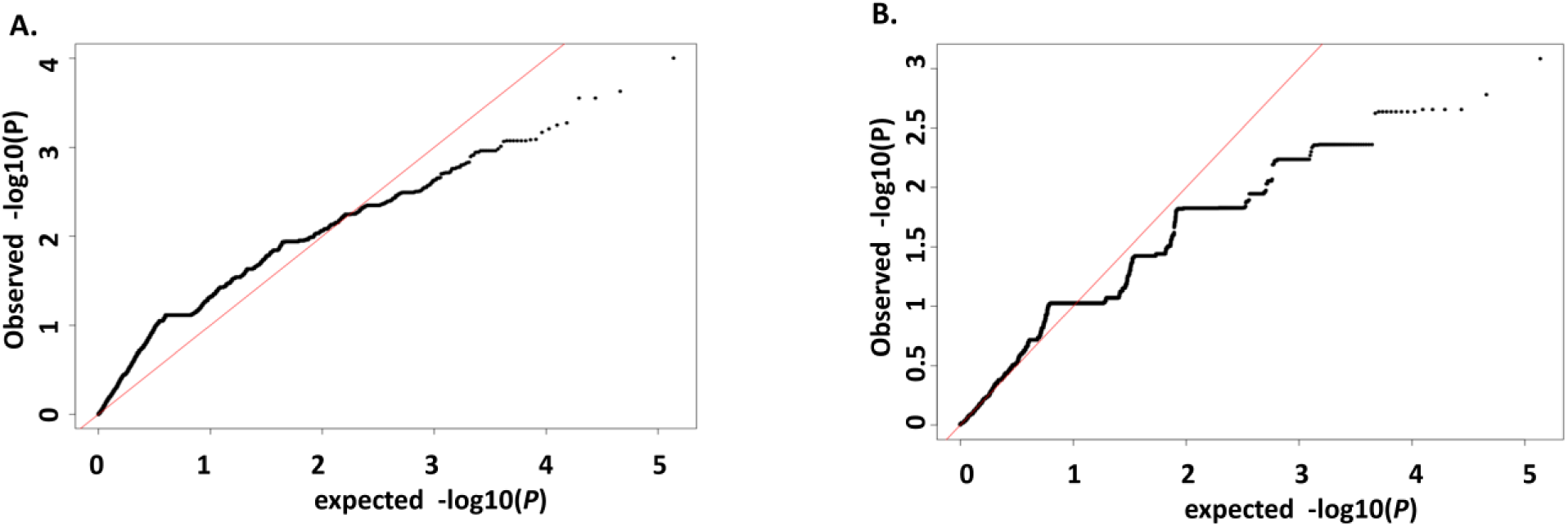
Quantile-quantile(Q-Q) plots of −log10 (*P-value*) from GWA analysis of *A. fumigatus* ITCZ sensitivity using Tassel (A) and RoadTrips (B). In each of the plots, Y-axis displays the quantile distribution of observed −log10 (*P-value*) while X-axis shows the quantile distribution of expected -log10 (*P-value*).

**Figure S4.**
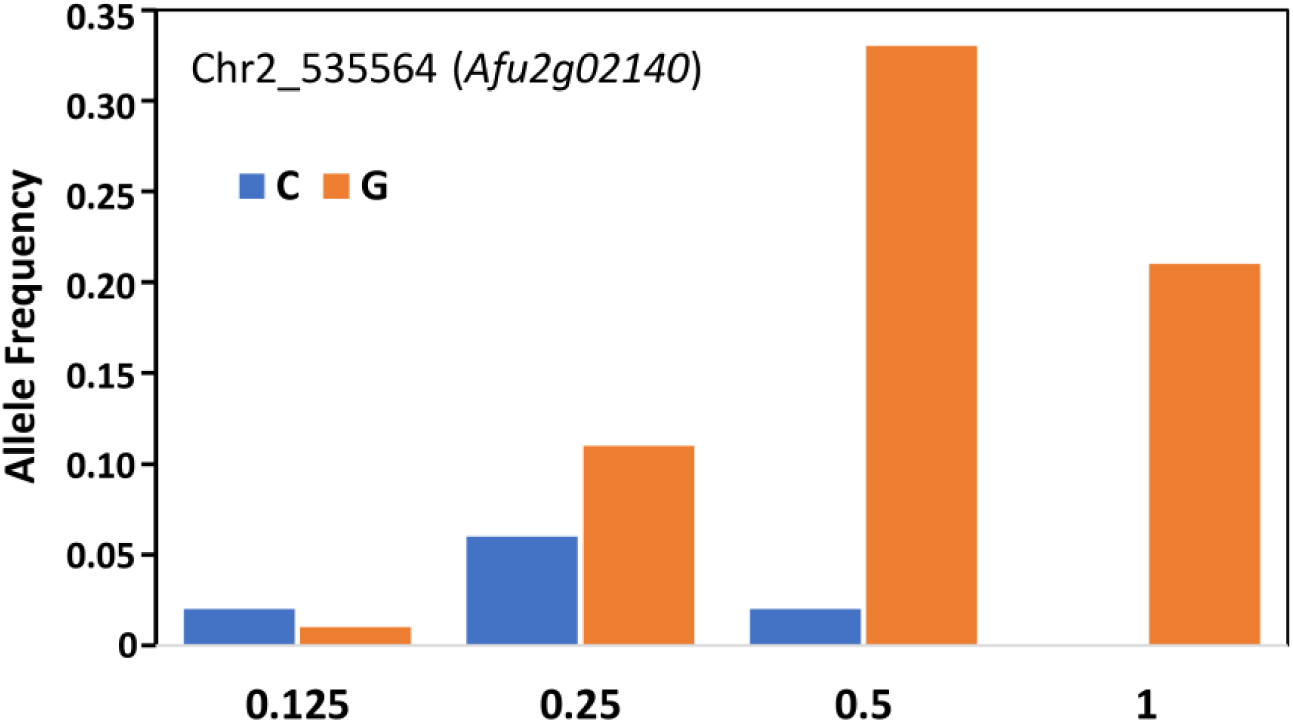
Allele frequency of the SNP in *Afu2g02140* that is associated with ITCZ sensitivity. The allele frequency of each allele is displayed on the Y-axis across different ITCZ MICs (X-axis). Blue and orange bars represent the minor and major alleles, respectively.

**Figure S5.**
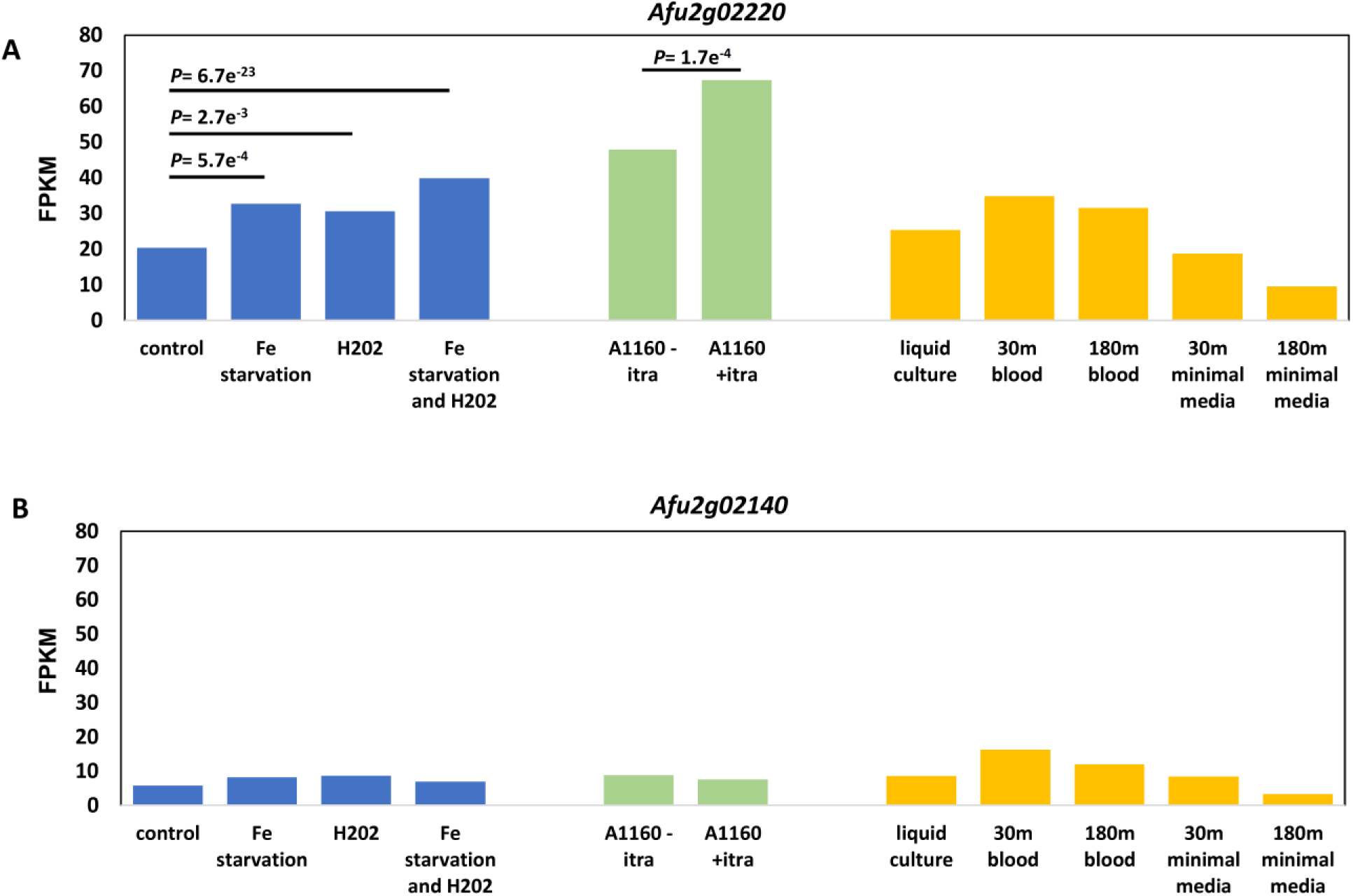
The expression of *Afu2g02220* (A) and *Afu2g02140* (B) in various conditions from RNA-seq data publicly available through FungiDB. FPKM (Fragments Per Kilobase of transcript per Million mapped reads) is displayed on Y-axis, while the X-axis represents experimental conditions. Bars are colored by study. *P*-values are reported for significant pairwise differential expression within the same study.

**Figure S6.**
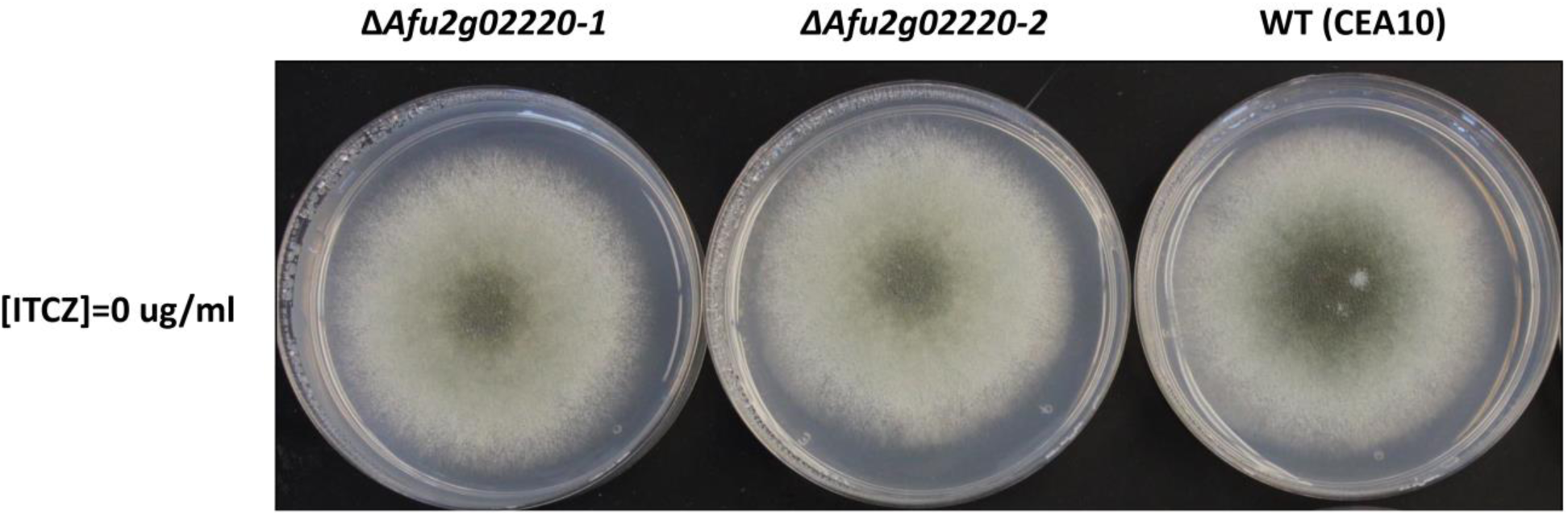
Growth of parent strain (CEA10) and *Afu2g02220* KOs on minimal media.

## REFERENCES

1. Brown GD, Denning DW, Gow NA, Levitz SM, Netea MG, White TC. 2012. Hidden killers: human fungal infections. Sci Transl Med 4:165rv13.

2. Latge JP. 1999. Aspergillus fumigatus and aspergillosis. Clin Microbiol Rev 12:310–50.

3. Latge JP, Chamilos G. 2019. Aspergillus fumigatus and Aspergillosis in 2019. Clin Microbiol Rev 33.

4. Neofytos D, Chatzis O, Nasioudis D, Boely Janke E, Doco Lecompte T, Garzoni C, Berger C, Cussini A, Boggian K, Khanna N, Manuel O, Mueller NJ, van Delden C, Swiss Transplant Cohort S. 2018. Epidemiology, risk factors and outcomes of invasive aspergillosis in solid organ transplant recipients in the Swiss Transplant Cohort Study. Transpl Infect Dis 20:e12898.

5. Robinett KS, Weiler B, Verceles AC. 2013. Invasive aspergillosis masquerading as catastrophic antiphospholipid syndrome. American Journal of Critical Care 22:448–451.

6. Lin SJ, Schranz J, Teutsch SM. 2001. Aspergillosis case-fatality rate: systematic review of the literature. Clin Infect Dis 32:358–66.

7. Latge JP, Beauvais A, Chamilos G. 2017. The Cell Wall of the Human Fungal Pathogen Aspergillus fumigatus: Biosynthesis, Organization, Immune Response, and Virulence. Annu Rev Microbiol 71:99–116.

8. Revie NM, Iyer KR, Robbins N, Cowen LE. 2018. Antifungal drug resistance: evolution, mechanisms and impact. Curr Opin Microbiol 45:70–76.

9. Alcazar-Fuoli L, Mellado E. 2013. Ergosterol biosynthesis in Aspergillus fumigatus: its relevance as an antifungal target and role in antifungal drug resistance. Frontiers in microbiology 3:439.

10. Garcia-Rubio R, Cuenca-Estrella M, Mellado E. 2017. Triazole Resistance in Aspergillus Species: An Emerging Problem. Drugs 77:599–613.

11. Fraczek MG, Bromley M, Bowyer P. 2011. An improved model of the Aspergillus fumigatus CYP51A protein. Antimicrob Agents Chemother 55:2483–6.

12. Warrilow AG, Parker JE, Price CL, Nes WD, Kelly SL, Kelly DE. 2015. In vitro biochemical study of CYP51-mediated azole resistance in Aspergillus fumigatus. Antimicrobial agents and chemotherapy 59:7771–7778.

13. Mellado E, Garcia-Effron G, Alcazar-Fuoli L, Melchers WJ, Verweij PE, Cuenca-Estrella M, Rodriguez-Tudela JL. 2007. A new Aspergillus fumigatus resistance mechanism conferring in vitro cross-resistance to azole antifungals involves a combination of cyp51A alterations. Antimicrob Agents Chemother 51:1897–904.

14. Hagiwara D, Watanabe A, Kamei K. 2016. Sensitisation of an Azole-Resistant Aspergillus fumigatus Strain containing the Cyp51A-Related Mutation by Deleting the SrbA Gene. Sci Rep 6:38833.

15. Camps SM, Dutilh BE, Arendrup MC, Rijs AJ, Snelders E, Huynen MA, Verweij PE, Melchers WJ. 2012. Discovery of a HapE mutation that causes azole resistance in Aspergillus fumigatus through whole genome sequencing and sexual crossing. PLoS One 7:e50034.

16. Paul S, Stamnes M, Thomas GH, Liu H, Hagiwara D, Gomi K, Filler SG, Moye-Rowley WS. 2019. AtrR is an essential determinant of azole resistance in Aspergillus fumigatus. MBio 10.

17. Fraczek MG, Bromley M, Buied A, Moore CB, Rajendran R, Rautemaa R, Ramage G, Denning DW, Bowyer P. 2013. The cdr1B efflux transporter is associated with non-cyp51a-mediated itraconazole resistance in Aspergillus fumigatus. Journal of Antimicrobial Chemotherapy 68:1486–1496.

18. Meneau I, Coste AT, Sanglard D. 2016. Identification of Aspergillus fumigatus multidrug transporter genes and their potential involvement in antifungal resistance. Sabouraudia 54:616–627.

19. Moye-Rowley W. 2015. Multiple mechanisms contribute to the development of clinically significant azole resistance in Aspergillus fumigatus. Frontiers in microbiology 6:70.

20. Chen P, Liu J, Zeng M, Sang H. 2020. Exploring the molecular mechanism of azole resistance in Aspergillus fumigatus. J Mycol Med 30:100915.

21. Hokken MWJ, Zoll J, Coolen JPM, Zwaan BJ, Verweij PE, Melchers WJG. 2019. Phenotypic plasticity and the evolution of azole resistance in Aspergillus fumigatus; an expression profile of clinical isolates upon exposure to itraconazole. BMC Genomics 20:28.

22. da Silva Ferreira ME, Malavazi I, Savoldi M, Brakhage AA, Goldman MH, Kim HS, Nierman WC, Goldman GH. 2006. Transcriptome analysis of Aspergillus fumigatus exposed to voriconazole. Curr Genet 50:32–44.

23. Gibson G. 2018. Population genetics and GWAS: a primer. PLoS biology 16:e2005485.

24. Chen PE, Shapiro BJ. 2015. The advent of genome-wide association studies for bacteria. Curr Opin Microbiol 25:17–24.

25. Power RA, Parkhill J, de Oliveira T. 2017. Microbial genome-wide association studies: lessons from human GWAS. Nat Rev Genet 18:41–50.

26. Read TD, Massey RC. 2014. Characterizing the genetic basis of bacterial phenotypes using genome-wide association studies: a new direction for bacteriology. Genome Med 6:109.

27. Dalman K, Himmelstrand K, Olson A, Lind M, Brandstrom-Durling M, Stenlid J. 2013. A genome-wide association study identifies genomic regions for virulence in the non-model organism Heterobasidion annosum s.s. PLoS One 8:e53525.

28. Muller LA, Lucas JE, Georgianna DR, McCusker JH. 2011. Genome-wide association analysis of clinical vs. nonclinical origin provides insights into Saccharomyces cerevisiae pathogenesis. Molecular ecology 20:4085–4097.

29. Gao Y, Liu Z, Faris JD, Richards J, Brueggeman RS, Li X, Oliver RP, McDonald BA, Friesen TL. 2016. Validation of genome-wide association studies as a tool to identify virulence factors in Parastagonospora nodorum. Phytopathology 106:1177–1185.

30. Palma-Guerrero J, Hall CR, Kowbel D, Welch J, Taylor JW, Brem RB, Glass NL. 2013. Genome wide association identifies novel loci involved in fungal communication. PLoS Genet 9:e1003669.

31. Talas F, Kalih R, Miedaner T, McDonald BA. 2016. Genome-Wide Association Study Identifies Novel Candidate Genes for Aggressiveness, Deoxynivalenol Production, and Azole Sensitivity in Natural Field Populations of Fusarium graminearum. Mol Plant Microbe Interact 29:417–30.

32. Hartmann FE, Sanchez-Vallet A, McDonald BA, Croll D. 2017. A fungal wheat pathogen evolved host specialization by extensive chromosomal rearrangements. ISME J 11:1189–1204.

33. Takahashi-Nakaguchi A, Muraosa Y, Hagiwara D, Sakai K, Toyotome T, Watanabe A, Kawamoto S, Kamei K, Gonoi T, Takahashi H. 2015. Genome sequence comparison of Aspergillus fumigatus strains isolated from patients with pulmonary aspergilloma and chronic necrotizing pulmonary aspergillosis. Med Mycol 53:353–60.

34. McKenna A, Hanna M, Banks E, Sivachenko A, Cibulskis K, Kernytsky A, Garimella K, Altshuler D, Gabriel S, Daly M, DePristo MA. 2010. The Genome Analysis Toolkit: a MapReduce framework for analyzing next-generation DNA sequencing data. Genome Res 20:1297–303.

35. Sul JH, Martin LS, Eskin E. 2018. Population structure in genetic studies: Confounding factors and mixed models. PLoS Genet 14:e1007309.

36. Alexander DH, Novembre J, Lange K. 2009. Fast model-based estimation of ancestry in unrelated individuals. Genome Res 19:1655–64.

37. Jombart T, Devillard S, Balloux F. 2010. Discriminant analysis of principal components: a new method for the analysis of genetically structured populations. BMC Genet 11:94.

38. Alexander DH, Lange K. 2011. Enhancements to the ADMIXTURE algorithm for individual ancestry estimation. BMC Bioinformatics 12:246.

39. Tashiro M, Izumikawa K, Minematsu A, Hirano K, Iwanaga N, Ide S, Mihara T, Hosogaya N, Takazono T, Morinaga Y. 2012. Antifungal susceptibilities of Aspergillus fumigatus clinical isolates obtained in Nagasaki, Japan. Antimicrobial agents and chemotherapy 56:584–587.

40. Price AL, Zaitlen NA, Reich D, Patterson N. 2010. New approaches to population stratification in genome-wide association studies. Nat Rev Genet 11:459–63.

41. Yu J, Pressoir G, Briggs WH, Bi IV, Yamasaki M, Doebley JF, McMullen MD, Gaut BS, Nielsen DM, Holland JB. 2006. A unified mixed-model method for association mapping that accounts for multiple levels of relatedness. Nature genetics 38:203–208.

42. Alam MT, Petit III RA, Crispell EK, Thornton TA, Conneely KN, Jiang Y, Satola SW, Read TD. 2014. Dissecting vancomycin-intermediate resistance in Staphylococcus aureus using genome-wide association. Genome biology and evolution 6:1174–1185.

43. Earle SG, Wu C-H, Charlesworth J, Stoesser N, Gordon NC, Walker TM, Spencer CC, Iqbal Z, Clifton DA, Hopkins KL. 2016. Identifying lineage effects when controlling for population structure improves power in bacterial association studies. Nature microbiology 1:1–8.

44. Bradbury PJ, Zhang Z, Kroon DE, Casstevens TM, Ramdoss Y, Buckler ES. 2007. TASSEL: software for association mapping of complex traits in diverse samples. Bioinformatics 23:2633–5.

45. Thornton T, McPeek MS. 2010. ROADTRIPS: case-control association testing with partially or completely unknown population and pedigree structure. The American Journal of Human Genetics 86:172–184.

46. Donaldson KM, Yin H, Gekakis N, Supek F, Joazeiro CA. 2003. Ubiquitin signals protein trafficking via interaction with a novel ubiquitin binding domain in the membrane fusion regulator, Vps9p. Curr Biol 13:258–62.

47. Shih SC, Prag G, Francis SA, Sutanto MA, Hurley JH, Hicke L. 2003. A ubiquitin-binding motif required for intramolecular monoubiquitylation, the CUE domain. EMBO J 22:1273–81.

48. Stajich JE, Harris T, Brunk BP, Brestelli J, Fischer S, Harb OS, Kissinger JC, Li W, Nayak V, Pinney DF, Stoeckert CJ, Jr., Roos DS. 2012. FungiDB: an integrated functional genomics database for fungi. Nucleic Acids Res 40:D675–81.

49. Kurucz V, Kruger T, Antal K, Dietl AM, Haas H, Pocsi I, Kniemeyer O, Emri T. 2018. Additional oxidative stress reroutes the global response of Aspergillus fumigatus to iron depletion. BMC Genomics 19:357.

50. Irmer H, Tarazona S, Sasse C, Olbermann P, Loeffler J, Krappmann S, Conesa A, Braus GH. 2015. RNAseq analysis of Aspergillus fumigatus in blood reveals a just wait and see resting stage behavior. BMC genomics 16:640.

51. Al Abdallah Q, Ge W, Fortwendel JR. 2017. A simple and universal system for gene manipulation in Aspergillus fumigatus: in vitro-assembled Cas9-guide RNA ribonucleoproteins coupled with microhomology repair templates. Msphere 2.

52. Hunt RC, Simhadri VL, Iandoli M, Sauna ZE, Kimchi-Sarfaty C. 2014. Exposing synonymous mutations. Trends Genet 30:308–21.

53. Warnecke D, Erdmann R, Fahl A, Hube B, Müller F, Zank T, Zähringer U, Heinz E. 1999. Cloning and Functional Expression of UGT Genes Encoding Sterol Glucosyltransferases from Saccharomyces cerevisiae, Candida albicans, Pichia pastoris, andDictyostelium discoideum. Journal of Biological Chemistry 274:13048–13059.

54. Gallego O, Betts MJ, Gvozdenovic-Jeremic J, Maeda K, Matetzki C, Aguilar-Gurrieri C, Beltran-Alvarez P, Bonn S, Fernández-Tornero C, Jensen LJ. 2010. A systematic screen for protein–lipid interactions in Saccharomyces cerevisiae. Molecular systems biology 6:430.

55. Kikuma T, Tadokoro T, Maruyama J-i, Kitamoto K. 2017. AoAtg26, a putative sterol glucosyltransferase, is required for autophagic degradation of peroxisomes, mitochondria, and nuclei in the filamentous fungus Aspergillus oryzae. Bioscience, Biotechnology, and Biochemistry 81:384–395.

56. Oku M, Warnecke D, Noda T, Muller F, Heinz E, Mukaiyama H, Kato N, Sakai Y. 2003. Peroxisome degradation requires catalytically active sterol glucosyltransferase with a GRAM domain. EMBO J 22:3231–41.

57. Asakura M, Ninomiya S, Sugimoto M, Oku M, Yamashita S-i, Okuno T, Sakai Y, Takano Y. 2009. Atg26-mediated pexophagy is required for host invasion by the plant pathogenic fungus Colletotrichum orbiculare. The Plant Cell 21:1291–1304.

58. Amarsaikhan N, Albrecht-Eckardt D, Sasse C, Braus GH, Ogel ZB, Kniemeyer O. 2017. Proteomic profiling of the antifungal drug response of Aspergillus fumigatus to voriconazole. Int J Med Microbiol 307:398–408.

59. Dunlap JC, Borkovich KA, Henn MR, Turner GE, Sachs MS, Glass NL, McCluskey K, Plamann M, Galagan JE, Birren BW, Weiss RL, Townsend JP, Loros JJ, Nelson MA, Lambreghts R, Colot HV, Park G, Collopy P, Ringelberg C, Crew C, Litvinkova L, DeCaprio D, Hood HM, Curilla S, Shi M, Crawford M, Koerhsen M, Montgomery P, Larson L, Pearson M, Kasuga T, Tian C, Basturkmen M, Altamirano L, Xu J. 2007. Enabling a community to dissect an organism: overview of the Neurospora functional genomics project. Adv Genet 57:49–96.

60. Colot HV, Park G, Turner GE, Ringelberg C, Crew CM, Litvinkova L, Weiss RL, Borkovich KA, Dunlap JC. 2006. A high-throughput gene knockout procedure for Neurospora reveals functions for multiple transcription factors. Proc Natl Acad Sci U S A 103:10352–10357.

61. Sardi M, Paithane V, Place M, Robinson E, Hose J, Wohlbach DJ, Gasch AP. 2018. Genome-wide association across Saccharomyces cerevisiae strains reveals substantial variation in underlying gene requirements for toxin tolerance. PLoS Genet 14:e1007217.

62. Leinonen R, Sugawara H, Shumway M, International Nucleotide Sequence Database C. 2011. The sequence read archive. Nucleic Acids Res 39:D19–21.

63. John H. 2008. Reference method for broth dilution antifungal susceptibility testing of filamentous fungi, approved standard. M38-A2. Clin Lab Stand Inst 28:1–35.

64. Zhao S, Latge JP, Gibbons JG. 2019. Genome Sequences of Two Strains of the Food Spoilage Mold Aspergillus fischeri. Microbiol Resour Announc 8.

65. Davis MP, van Dongen S, Abreu-Goodger C, Bartonicek N, Enright AJ. 2013. Kraken: a set of tools for quality control and analysis of high-throughput sequence data. Methods 63:41–9.

66. Nierman WC, Pain A, Anderson MJ, Wortman JR, Kim HS, Arroyo J, Berriman M, Abe K, Archer DB, Bermejo C, Bennett J, Bowyer P, Chen D, Collins M, Coulsen R, Davies R, Dyer PS, Farman M, Fedorova N, Fedorova N, Feldblyum TV, Fischer R, Fosker N, Fraser A, Garcia JL, Garcia MJ, Goble A, Goldman GH, Gomi K, Griffith-Jones S, Gwilliam R, Haas B, Haas H, Harris D, Horiuchi H, Huang J, Humphray S, Jimenez J, Keller N, Khouri H, Kitamoto K, Kobayashi T, Konzack S, Kulkarni R, Kumagai T, Lafon A, Latge JP, Li W, Lord A, Lu C, et al. 2005. Genomic sequence of the pathogenic and allergenic filamentous fungus Aspergillus fumigatus. Nature 438:1151–6.

67. Li H, Durbin R. 2009. Fast and accurate short read alignment with Burrows-Wheeler transform. Bioinformatics 25:1754–60.

68. Li H, Handsaker B, Wysoker A, Fennell T, Ruan J, Homer N, Marth G, Abecasis G, Durbin R, Genome Project Data Processing S. 2009. The Sequence Alignment/Map format and SAMtools. Bioinformatics 25:2078–9.

69. Van der Auwera GA, Carneiro MO, Hartl C, Poplin R, Del Angel G, Levy-Moonshine A, Jordan T, Shakir K, Roazen D, Thibault J, Banks E, Garimella KV, Altshuler D, Gabriel S, DePristo MA. 2013. From FastQ data to high confidence variant calls: the Genome Analysis Toolkit best practices pipeline. Curr Protoc Bioinformatics 43:11 10 1–11 10 33.

70. Danecek P, Auton A, Abecasis G, Albers CA, Banks E, DePristo MA, Handsaker RE, Lunter G, Marth GT, Sherry ST, McVean G, Durbin R, Genomes Project Analysis G. 2011. The variant call format and VCFtools. Bioinformatics 27:2156–8.

71. Jombart T. 2008. adegenet: a R package for the multivariate analysis of genetic markers. Bioinformatics 24:1403–5.

72. Team RC. 2013. R: A language and environment for statistical computing. Vienna, Austria.

73. Huson DH, Bryant D. 2006. Application of phylogenetic networks in evolutionary studies. Molecular biology and evolution 23:254–267.

74. Turner SD. 2014. qqman: an R package for visualizing GWAS results using QQ and manhattan plots. Biorxiv:005165.

75. Cingolani P, Platts A, Wang le L, Coon M, Nguyen T, Wang L, Land SJ, Lu X, Ruden DM. 2012. A program for annotating and predicting the effects of single nucleotide polymorphisms, SnpEff: SNPs in the genome of Drosophila melanogaster strain w1118; iso-2; iso-3. Fly (Austin) 6:80–92.

76. Peng D, Tarleton R. 2015. EuPaGDT: a web tool tailored to design CRISPR guide RNAs for eukaryotic pathogens. Microb Genom 1:e000033.

77. Shimizu K, Keller NP. 2001. Genetic involvement of a cAMP-dependent protein kinase in a G protein signaling pathway regulating morphological and chemical transitions in Aspergillus nidulans. Genetics 157:591–600.

